# ‘Macrobot’–an automated segmentation-based system for powdery mildew disease quantification

**DOI:** 10.1101/2020.03.16.993451

**Authors:** Stefanie Lück, Marc Strickert, Maximilian Lorbeer, Friedrich Melchert, Andreas Backhaus, David Kilias, Udo Seiffert, Patrick Schweizer, Armin Djamei, Dimitar Douchkov

**Author notes:** Deceased 09.03.2018 (Patrick Schweizer).

## Abstract

Plant diseases, as one of the perpetual problems in agriculture, is increasingly difficult to manage due to intensifying of the field production, global trafficking, reduction of genetic variability of crops, climatic changes-driven expansion of pests, redraw and loss of effectiveness of pesticides and rapid breakdown of the disease resistance in the field. The substantial progress in genomics of both plants and pathogens, achieved in the last decades has the potential to counteract this negative trend, however, only when the genomic data is supported by relevant phenotypic data that allows linking the genomic information to specific traits. In this respect, phenotyping is and will remain an essential element of any comprehensive functional genomics study.

We have developed a set of methods and equipment and combined them into a “Macrophenomics pipeline”. The pipeline has been optimized for the quantification of powdery mildew infection symptoms on wheat and barley but it can be adapted to other diseases and host plants. The Macrophenomics pipeline scores the visible disease symptoms, typically 5-7 days after inoculation (dai) in a highly automated manner. The system can precisely and reproducibly quantify the percentage of the infected leaf area with a throughput of the image acquisition module of up to 10 000 individual samples per day, making it appropriate for phenotyping of large germplasms collections and crossing populations.

## 1. Introduction

Cereals, which include wheat, barley, rice, maize, rye, oats, sorghum, and millet, have been the primary component of humans diet delivering more than 50% of the world’s daily caloric intake(Awika 2011). Like any other plant, these species are under constant attack by a vast number of pathogens. Since the impact of cereal diseases is proportional to the importance of these crops for human nutrition they are of exceptional interest to plant pathologists and breeders.

Precise and sensitive phenotyping is one of the key requirements for modern breeding approaches and functional genomics studies. Many of the desired traits and phenotypes are polygenic by nature and their manifestation depends on the cumulative effect of several factors with small to moderate effect. The quantitative disease resistance of the plants against pathogens is a typical example of a complex polygenic trait. Although this type of resistance is usually less efficient than the strong R-gene based resistance, it is nevertheless desired by the breeders because it is more durable on the field and in great contrast to the R-gene resistance, it is effective against all races of a particular pathogen and even against different pathogen species. But studying the underlying mechanisms of the quantitative resistance is seriously challenged by the complexity of this phenomenon (Corwin and Kliebenstein 2017, Jones and Dangl 2006). The accessibility of the genomic information for several host and pathogen species greatly facilitates these studies but on the other hand, introduced an enormous amount of data that needs to be tested and functionally validated. Thus, the ability of high-throughput become a major requirement for the new systematic phenotyping, and the term “phenomics” was coined to describe this approach.

The natural disease resistance is, beside the high yield and abiotic stress resistance, one of the most desired traits since the beginning of the agriculture. The breeders invested significant efforts in improving these traits and as a result, the modern crop cultivars are usually outperforming their wild progenitors in nearly all aspects. But unlike other factors that may influence plant performance, the pathogens constantly develop and modify strategies to evade the host defense mechanisms in a process some times called “evolutionary arms race”.

Powdery mildew (PM) is a disease caused by a diverse group of obligate biotrophic fungi that lead to extensive damage on various crop plants including cereals. *Blumeria graminis* is the causative agent of the powdery mildew disease of wheat and barley (Bockus et al. 2010). As most of the obligate biotrophs, *B. graminis* shows extreme host specificity. So-called *formae speciales* (f.sp.) have specialized virulence for particular cereals, e.g. for barley (*B. graminis* f. sp. hordei) or wheat, (*B. graminis* f.sp. tritici).

The asexual life cycle of *B. graminis* is fast and completes within a week. The haploid asexual fungal spores, called conidia, start germination within a few hours after contact with a plant leaf. The appressorial germ tube penetrates directly the cell wall of the leaf epidermal cells and grows into the living plant cell forming a feeding structure called haustorium. The establishment of biotrophy occurs within the first 24 hours after leaf spore inoculation. In the following days epiphytically growing hyphae develop many secondary haustoria in neighboring epidermal cells next to the initial infection site. After 3 days the fungal colony is macroscopically visible. In the following days, abundant spores are formed by the mycelium which completes the life cycle (Jankovics et al. 2015). In controlled infection assays with defined spore titters, the severity of infection and the size of the infected area is commonly the scoring parameter in disease rating to estimate host susceptibility (Nicot et al. 2002).

The common wheat (*Triticum aestivum*) is one of the most important staple food worldwide (Gennari and Monvayo 2018). Cultivated barley (*Hordeum vulgares* spp. *vulgare*), which is another member of the same *Triticeae* tribe, is also among the most favored crops worldwide. Besides, barley is an important genetic model for the very closely related but more complex wheat genome.

With the significant progress made on the sequencing of several cereal genomes and those of the corresponding powdery mildews, Genome Wide Association Studies (GWAS) to identify resistance traits became possible. However, a bottleneck for successful genotype-phenotype associations is the high throughput monitoring of disease symptom development as a measure of host plant susceptibility. Disease resistance traits range from partial, or quantitative, to complete, or qualitative. It has been shown in many cases that quantitative disease resistance is more durable on the field and therefore, of high potential value to the breeders (Johnson 1981). However, the quantitative resistance is usually a polygenic trait, which is based on the joined effect of many genes, where each of them contributes quantitatively to the level of plant defense (Niks, Qi, and Marcel 2015). The identification of genes with small to moderate resistance effects requires very precise and reproducible quantification of infection as a pre-requisite for genetic fine mapping and gene isolation. We have developed a set of methods and equipment and combined them into a “Macrophenomics pipeline”. The pipeline has been optimized for the quantification of Powdery mildew infection symptoms on wheat and barley but it can be adapted to other diseases and host plants.

## 2. Materials and Methods

### 2.1. Experimental design

The Macrophenomics pipeline consists of hard- and software components. A specialized robotic system implements the image acquisition part of the Macrophenomics pipeline. The so-called Macrobot autonomously acquires images of detached leaf segments mounted on standard size microtiter plates (MTP) filled with 1% water-agar for keeping the humidity (**Figure 1**).

**Figure 1.**
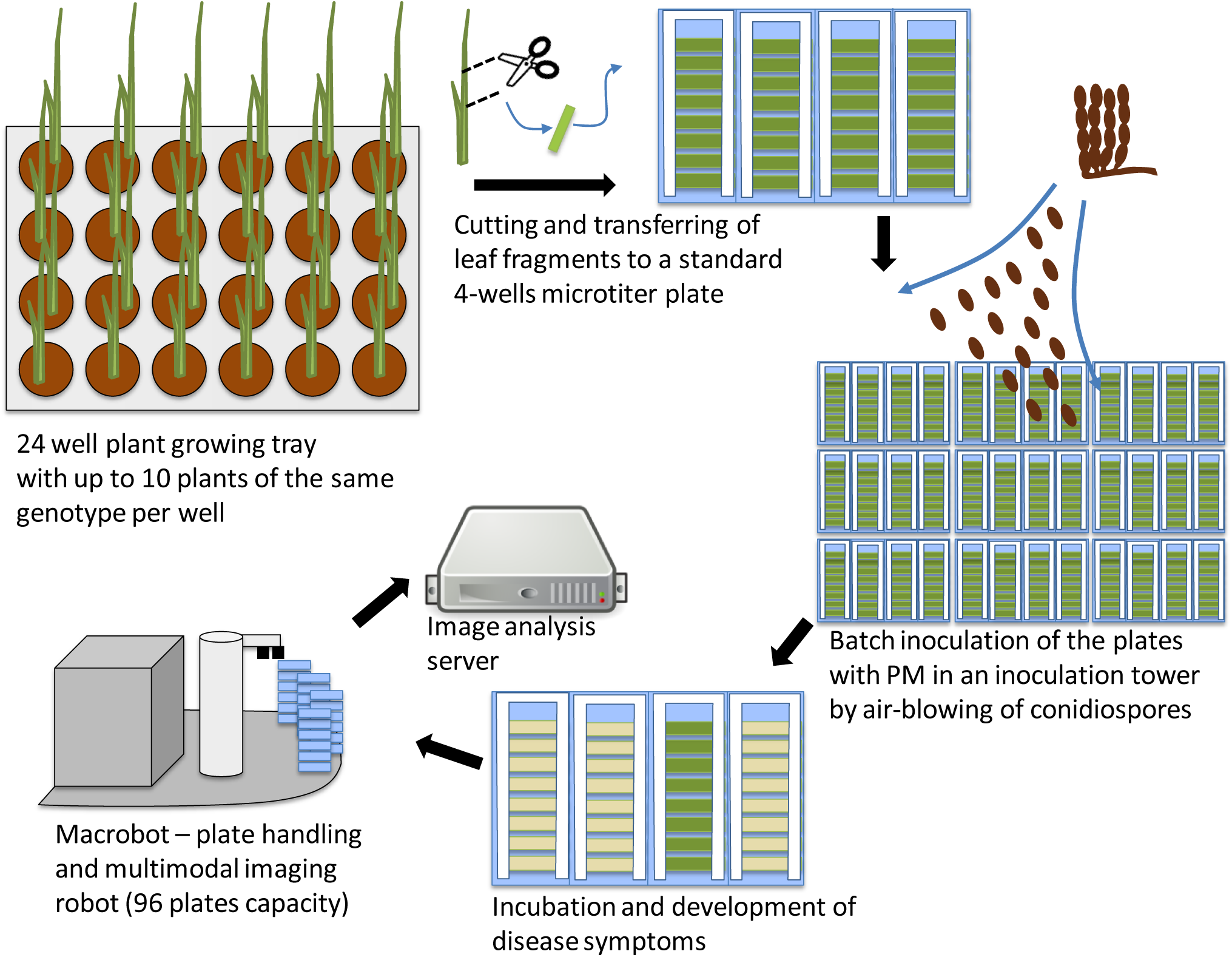
Overview of the phenotyping pipeline. The plants are grown in 24-well trays in a greenhouse. At appropriate stage, leaf fragments are harvested and mounted on standard 4-well microtiter plates and inoculated by air-blowing of powdery mildew spores in an inoculation tower. After incubation of 5-7 days, depending on the experiment design, the disease symptoms are clearly visible. The plates with the infected leaves are loaded into the Macrobot system for automated imaging. The acquired images are transferred to the image analysis server for quantification of the disease symptoms.

Typically, the wheat and barley plants are grown in 24-well trays in a greenhouse. The samples are taken at the 2-leaf stage from the middle part of the second leaf. The leaf fragments are mounted on standard 4-well MTPs with 1% water agar (Phyto agar, Duchefa, Haarlem, The Netherlands) supplemented by 20 mg.L^-1^ benzimidazole as a leaf senescence inhibitor. To achieve equal inoculation of all leaves, the plates are placed without lids in a rotating table inside an inoculation tower and are inoculated by blow-in of conidiospores from sporulating material. Inoculated plates are incubated in environmentally controlled plant growth chambers (20°C, 60%RH constant; 16 h light, 15 µE m^-2^ s^-1^) for 6 days until the disease symptoms are visible. The infected plates are loaded into the Macrobot system for automated imaging. The acquired images are transferred to the image analysis server for quantification of the disease symptoms.

### 2.2. Hardware

In the original version, the Macrobot employs a 14-bit monochrome camera (Thorlabs 8050M-GE-TE) at a resolution of 3296×2472 px. A high-end lens (CoastalOpt UV-VIS-IR 60 mm 1:4 Apo Macro) with apochromatic correction in the range from 310 to 1100 nm wavelength ensures that images using different illumination setups are precisely registered and focused. The illumination is realized using small bandwidth isotropic LED light sources (Metaphase Exolight-ISO-14-XXX-U) with 365nm (UV), 470nm (blue), 530nm (green) and 625nm (red) peak wavelength.

For each plate monochrome images in all illumination wavelengths are acquired separately and stored in 16-bit TIFF image files. An RGB image is generated by combining the images of the red, green and the blue LED channels (Supplemental figure S1). The UV channel is used to facilitate the extraction of the region of interest (ROI), where the leaves are located.

An improved version of Macrobot was introduced on a later stage and designated as Macrobot 2.0 (**Figure 2**). The illumination system was upgraded by doubling the LED units allowing bilateral illumination of the objects. A background illumination system based on electroluminescence foil was mounted on the MTP carrier to simplify the separation between the foreground and background, thus improving the leaf segmentation. The image acquisition and hardware controlling software was upgraded to 64-bit version for optimal system memory utilization. The entire technical layout was improved in respect of the gained experience with the first version of the Macrobot. Since the image acquisition components remains basically unchanged, data generated by Macrobot 2.0 is fully comparable to data acquired by the Macrobot 1.0, as far as compatible hardware setup is used. The data presented in this article was acquired by the original Macrobot hardware configuration.

**Figure 2.**
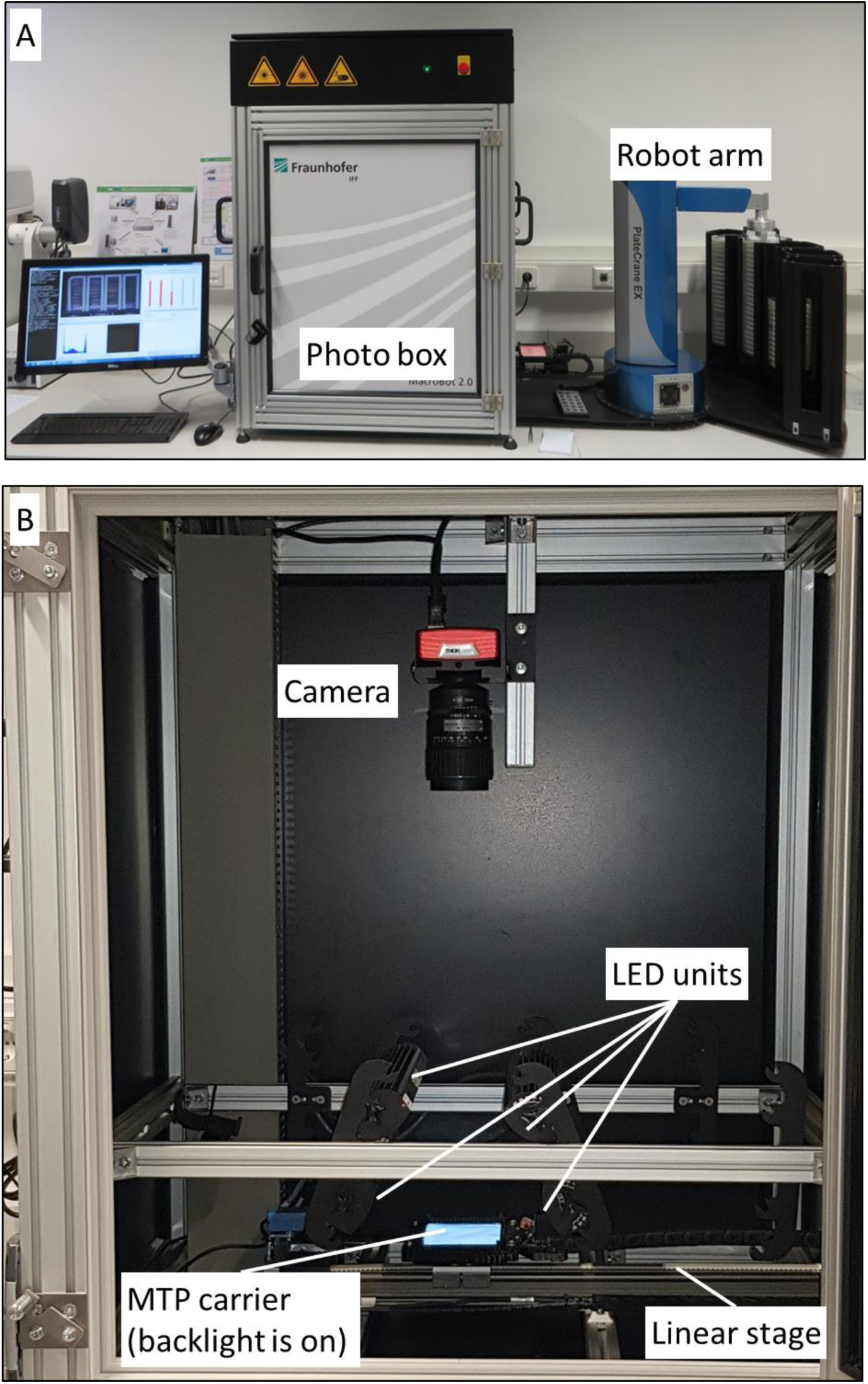
Macrobot 2.0 with improved technical design, bilateral illumination and background light. A - Outside view; B - Inside view of the photo box.

### 2.3. Software

The software was implemented in Python 2.7 under Windows 7 with extensive use of the NumPy (v. 1.12.1) (Walt, Colbert, and Varoquaux 2011), opencv-python (v. 2.4.13), scikit-learn (v. 0.17.1) (Pedregosa et al. 2011) and scikit-image (v. 0.13.0) (Walt, Colbert, and Varoquaux 2011) open source libraries. The source code is available at (Lueck 2019).

### 2.4. Model evaluation

Each model was validated by calculating the accuracy, recall, and precision of the model to test the prediction performance for each class. The overall accuracy is calculated by the number of correctly predicted observations divided by the total number of observations:

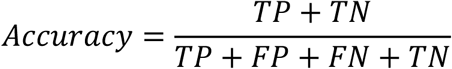

The precision is a measure of the false positive rate. It can be calculated by dividing the true positive observations by the total predicted positive observations:

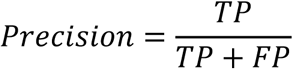

The recall measures the sensitivity of the predicted positive observations:

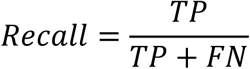

### 2.5. Plant and fungal material

Wheat and barley plants from different cultivars and landraces were grown in 24-pots trays (31×53 cm) in a greenhouse at 20°C constant, 16 h light period in a soil substrate. The first or the second leaves were harvested at 7 days, resp. 13-14 days after sowing. The leaf segments were mounted on 20 mg.L^-1^ benzimidazole supplemented, 1% water agar plates and inoculated with the corresponding pathogen at approximately 10 spores/mm^2^. As pathogens, the Swiss wheat powdery mildew field isolate FAL 92315 and the Swiss barley powdery mildew field isolate CH4.8, were used respectively. The image acquisition was performed seven days after inoculation (dai).

### 2.6. Quantitative PCR

Quantitative real-time PCR was performed in a volume of 5 mL QuantiTect Probe PCR Kit (Qiagen GmbH, Hilden, Germany) kit and an ABI 7900HT fast real-time PCR system (ThermoFisher Scientific Inc., Waltham, MA, USA). Forty cycles (15 sec. 94°C, 30 sec. 56°C, 30 sec. 72°C, preceded by standard denaturation steps at 94°C for 2 min.) were conducted. Data were analyzed by the Standard curve method using the SDS 2.2.1 software (ThermoFisher Scientific Inc., Waltham, MA, USA). Standard curve dilution series were included for each gene, as fivefold dilutions and three technical replicates per DNA sample. The detected quantity of the fungal gene GTFI (beta-1,3-glucanosyltransferase, GenBank: EU646133.1) was normalized to the quantity of the barley UBC gene (Ubiquitin-conjugating enzyme, GenBank: AY220735.1) and used as a proxy for fungal biomass. Used primers and probes: for the powdery mildew GTFI gene - BgGTF1_F (5’TTGGCCAAACAACTCAACTC3’), BgGTF1_R (AGCAGACCAAGACACACCAG) and BgGTF1_PR (fluorescent TaqMan probe, FAM-5’CTCCCAGCAACACTCCAGCT3’-BHQ1); for the barley UBC gene - HvUBC_F (5’ACTCCGAAGCAGCCAGAATG3’); HvUBC_R (5’GATCAAGCACAGGGACACAAC3’) and HvUBC_PR (fluorescent TaqMan probe Yakima Yellow-5’GAGAACAAGCGCGAGTACAACCGCAAGGTG3’-BHQ1).

## 3. Results

### 3.1. Frame and leaf segmentation

To define the area where the leaf segments are located on the plates the C-shaped white frames that hold the leaves were segmented and extracted. Optimal results were achieved by applying a Otsu’s thresholding (Otsu 1979) on the UV image, followed by dilation with 8×8 kernel to obtain a binary image (**Figure 3**). A Moore-Neighbour (Weisstein 2019) tracing was used to extract the contours of the binary image and filter the frames by size and position.

**Figure 3.**
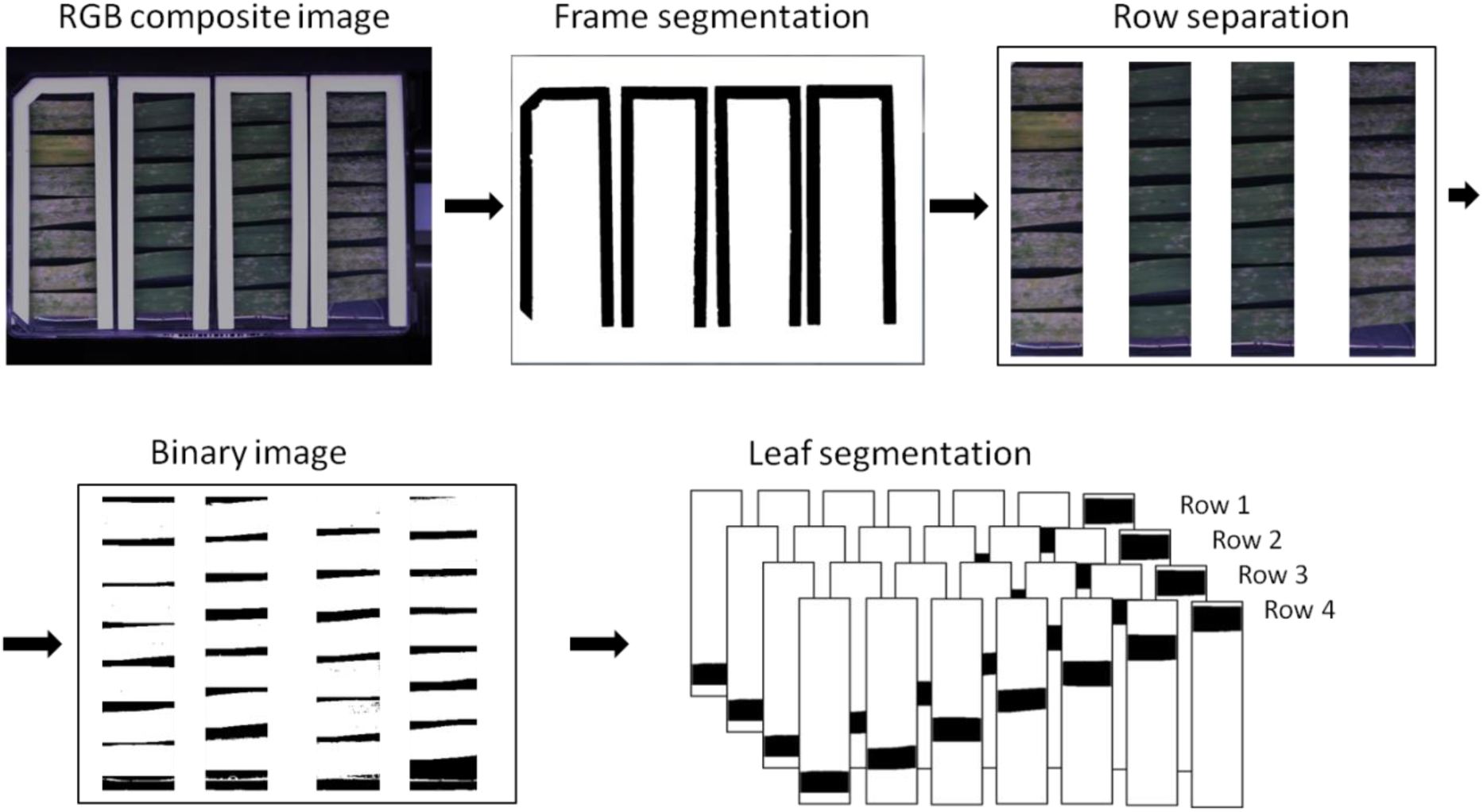
Frame and leaf segmentation processing chain

Each leaf segment was extracted to a separate region of interest (ROI). Best segmentation results were obtained by the Otsu’s binarization method on the backlight image, followed by Moore-Neighbour contour finding algorithm and object size selection. Otsu’s method also dealt with the particular challenge of interrupted leaf contours caused by necrosis or fungal infections.

### 3.2. Machine learning approach

The application of Machine learning approaches gives the advantage of using a data-driven analysis rather than hypothesis-driven statistics. In this way, complex statistical modelling assumptions can be reduced, offering possibly meaningful data features from which machine learning tools can derive desired classification outcomes in the manner of teaching. Therefore, several Machine learning methods were implemented and evaluated for their accuracy and performance in the quantification of the PM disease symptoms.

#### 3.2.1. Training data

Training data was collected by manual labelling of background, infected and leaf necrosis areas. The labelled single pixels were extracted and assigned to these three classes. To avoid a class imbalance the number of training samples per class was adjusted to the lowest number of pixels per class, which was 5 000. The dataset was split 70 % for training and 30 % for validation.

#### 3.2.2. Feature extraction and classification

We have compared three common classifiers - C-Support Vector Classification (Novakovic J 2011), Linear Support Vector Classification (Cortes and Vapnik 1995), and Random Forest (Tin Kam 1995). We found Random Forest as performing significantly better than Linear Support Vector Classification and slightly better compared to the C-Support Vector Classification (**Figure 4A, Supplemental table S2**). The training time with the Random Forest classifier was about 10 times faster as with the C-Support Vector Classification, therefore we end up using the Random Forest classifier for further experiments

**Figure 4.**
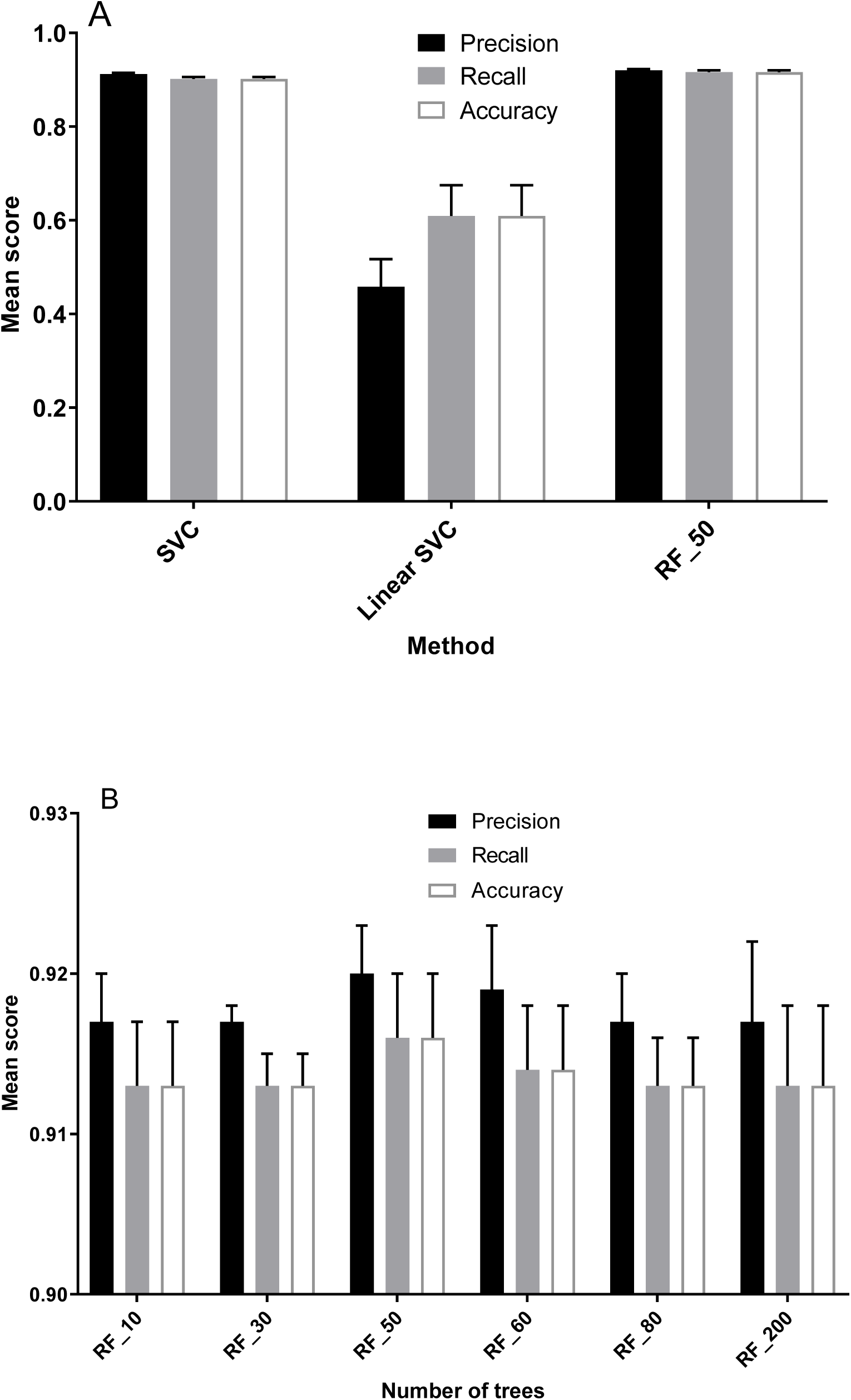
**A** - Evaluation of different classifiers on HSV_H_channel (5000 pixels/class; n=10; error bars=SD). **B** - evaluation of Random forest classifier with different numbers of trees on HSV_H_channel (5000 pixels per class, n=10).

To find the optimal number of trees for the Random Forest classifier, we tested six different values ranging from 10 to 200 trees, which lead to an optimum of 50 trees (**Figure 4B, Supplemental table S3**).

A Random Forest classifier has been trained by using RGB, LAB, HSV as multiple as well as single color channels (**Figure 5A, Supplemental table S4**).

**Figure 5.**
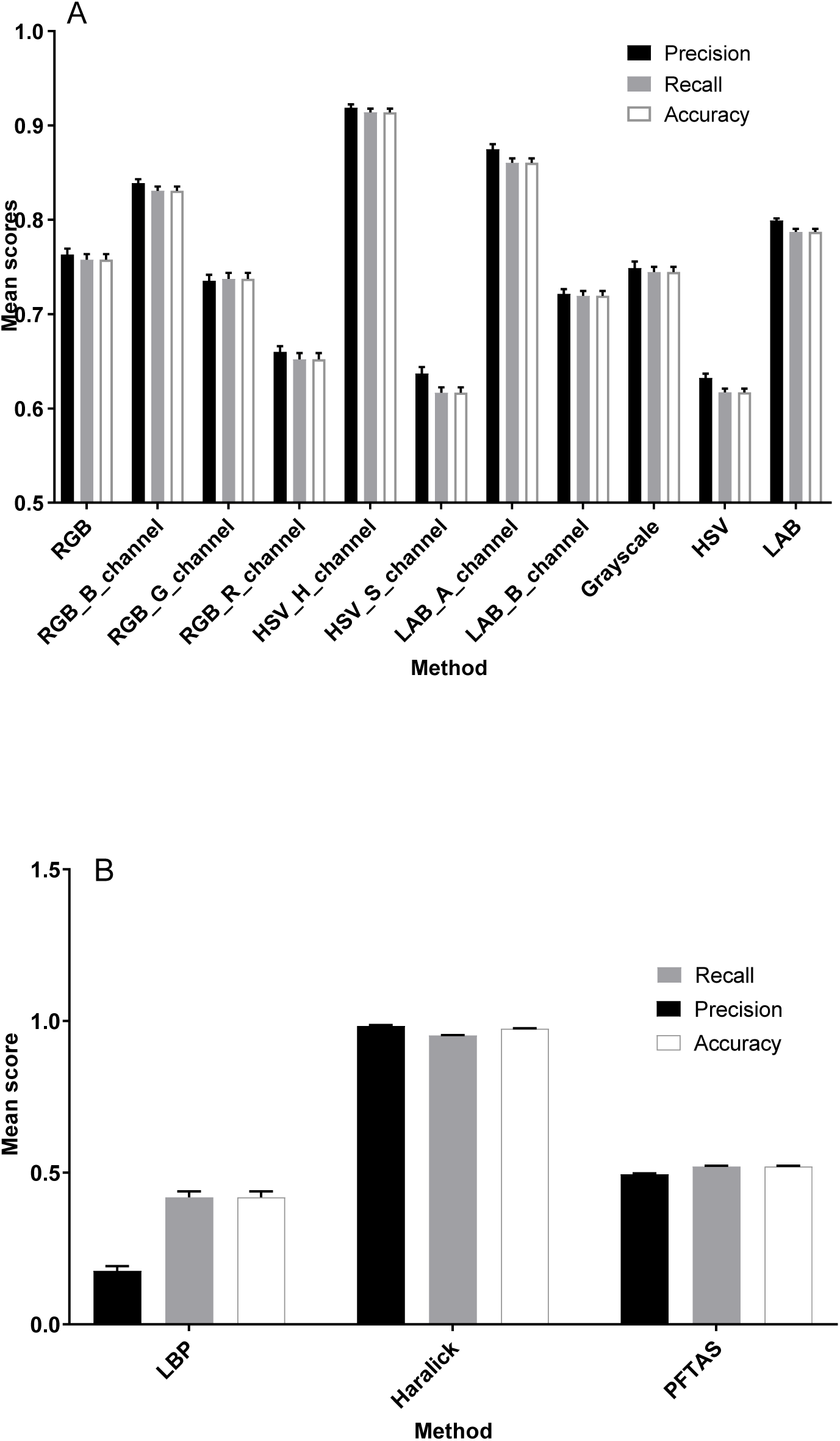
**A** - Evaluation of different color pixel classification method (5000 pixels/class; n=10; error bars=SD); Random Forest classifier (nr_trees=50). **B** - Evaluation of texture features (5000 pixels/class; n=10; error bars=SD); Random Forest classifier (nr_trees=50)

Texture spatial features as Local Binary Pattern (Ojala, Pietikainen, and Harwood 1994), Haralick (Haralick, Shanmugam, and Dinstein 1973) and Parameter Free Threshold Adjacency Statistics (Coelho et al. 2010) were also tested for improving the performance of the classifier. (**Figure 5B, Supplemental table S5**).

Four models reached an overall accuracy above 0.80, the blue channel of the RGB color space, the Hue channel of the HSV color space, the *a* channel of the *Lab* color space and the Haralick texture features. Those models were tested further in the validation experiment.

### 3.3. Segmentation approach

In addition to the Machine learning approach, we have tested several segmentation methods - Edge detection, Superpixel segmentation, Watershed transformation, Region-growing methods, Thresholding, Minimum and Maximum RGB (data not shown). The most efficient segmentation was achieved by the relatively simple method of Minimum RGB (minRGB) (**Figure 6**). The algorithm takes the single values for each RGB channel, determines the minimum number of each channel and stores the value. The other two channels are set to the value 0. This simple filter allowed reliable differentiation of the disease symptoms from the background by simultaneous reduction of the analysis artefacts and hardware workload.

**Figure 6.**
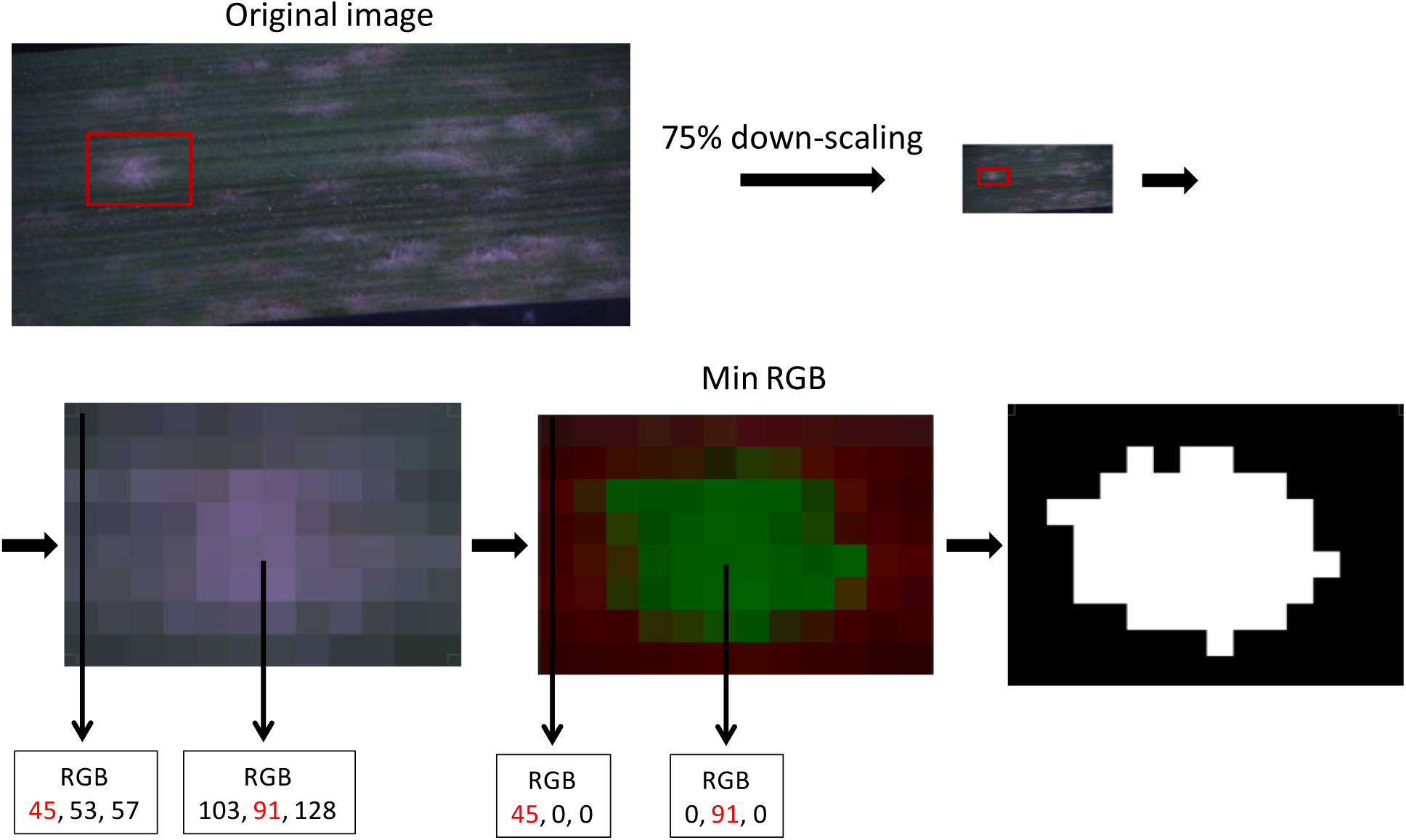
Minimal RGB value approach for segmentation.

### 3.4. Validation experiment

The Macrophenomics module aims to provide a precise and reproducible evaluation of the experimental results, and at the same time to release the human personnel from a routine and laborious task. However, over the years, a vast amount of manually obtained phenotypic data for many cultivars and landraces was accumulated and classified into common standards for disease rating. An automatic disease rating system that generates results in a good consistency with this data would be of particular interest to both plant pathologists and breeders. In this study, we have tested several computational methods and prediction models and we have selected the best performing. Still, it is not uncommon even for very accurate models to provide unsatisfactory results in the praxis. To estimate the performance of the different approaches and computer models on real-life data, we have carried out a validation experiment, where six experts were asked to do manual disease rating of the validation material. Combining the scores given by all experts formed a robust mean value, which was used to validate the computer prediction results. The validation set included a partially very difficult to score material with a lot of leaf senescence and necrosis.

In parallel to the visual methods, two other types of measurements were included for comparison – quantification of total fungal biomass using quantitative real-time PCR (qPCR) of fungal DNA, and inoculum density as number of applied fungal spores per mm^2^ of leaf surface (**Figure 9, Supplemental table S1**).

**Figure 7.**
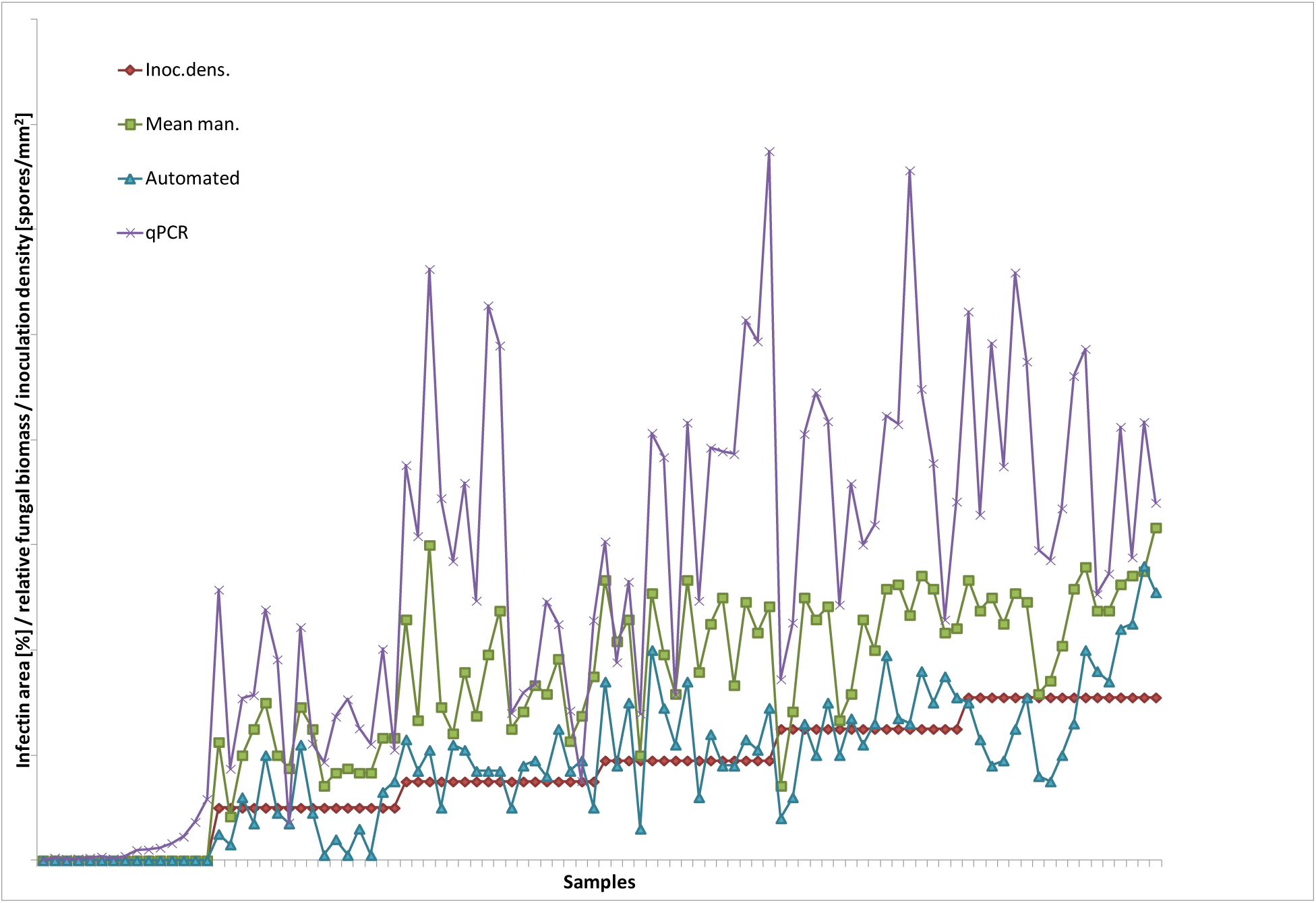
Plots of infection area determined automatically (blue triangles), mean manual values (“Mean man.”, green rectangles), together with the fungal biomass measured by qPCR (normalized relative transcript levels multiplied by 100 for better visibility; purple crosses) and inoculation density (spores per mm^2^, red rectangle, sorted ascending). On X-axis are ordered the samples, on Y-axis are the infection area (% of the leaf surface), resp. relative fungal biomass (relative units) and inoculation density (spores/mm^2^).

**Figure 8.**
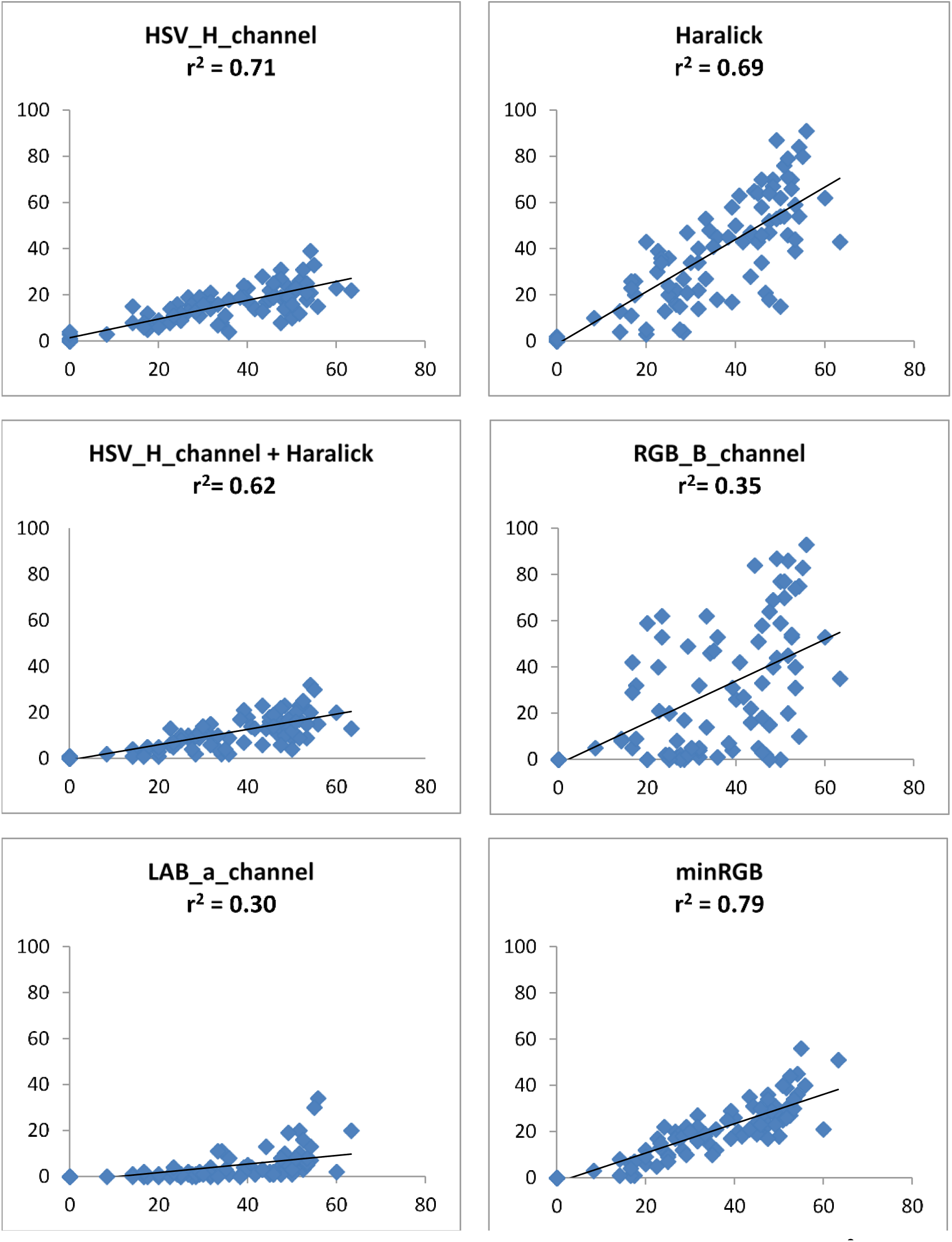
Pearson correlation plots and coefficients of determination (r^2^) for the different machine learning models and the minRGB based segmentation. On the horizontal axis are the mean manual scores and on the vertical axis are the algorithm prediction results. PM infected detached barley leaves, 6 dai. Number of samples n = 108, manual data is formed as a mean score of six testers.

**Figure 9.**
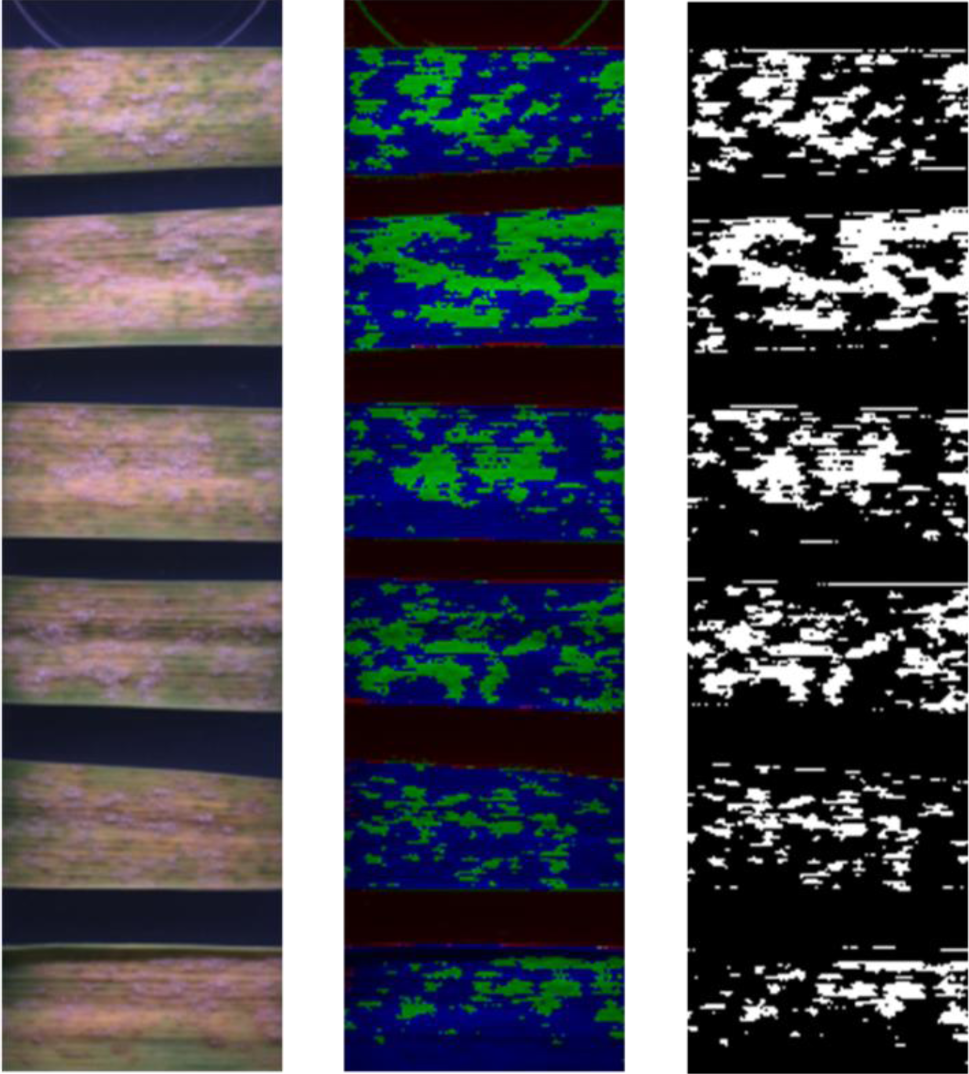
Example of a minRGB filter-based detection of disease area on PM infected detached barley leaves 6 dai. From left to right: RGB composite image, minRGB filter results, final prediction after thresholding.

#### 3.4.1. Performance of the Machine learning approach

All machine learning models with accuracy above 0.8 (**Figure 4** and **Figure 5**) plus minRGB segmentation algorithm were tested on validation experiment data. The prediction accuracy measured as Pearson correlation to the mean manual scores is shown in **Figure 8**.

#### 3.4.2. Performance of the segmentation approach

Although the leaf material of the validation experiment was often covered by large necrotic and/or chlorotic areas, which may complicate the disease recognition even for experienced laboratory personnel, the minRGB based prediction was very accurate (**Figure 9**).

The minRGB based algorithm was tested also in a large experiment with wheat, showing an even higher level of accuracy (**Figure 10**). The better results for the wheat material might be explained with the lower frequency of appearance of problematic artefacts as necrosis and senescence in this particular material.

**Figure 10.**
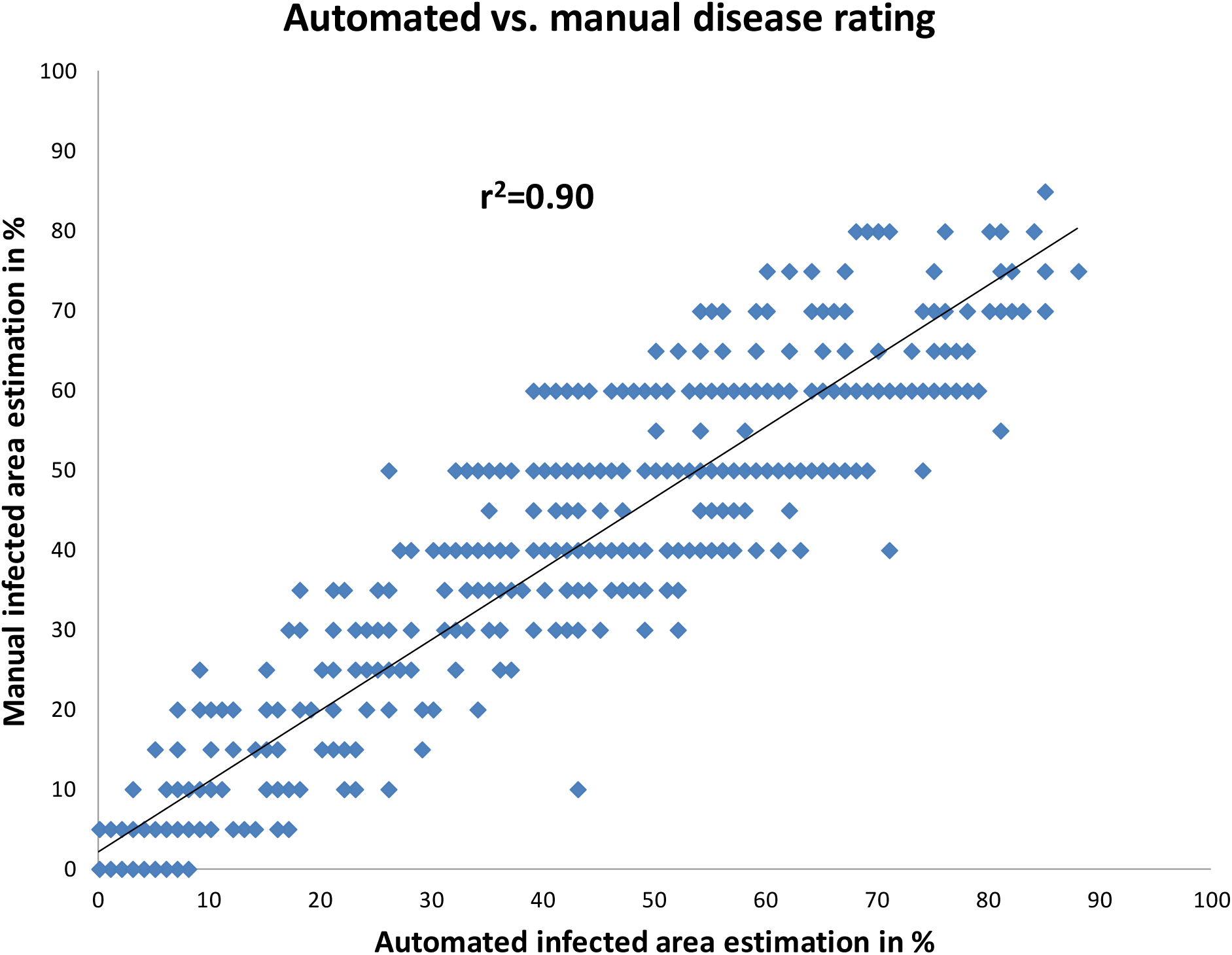
Correlation between the manual and minRGB based automated scores for PM infected detached wheat leaves, 6 dai. Number of samples n = 660, number of testers p = 1. The manual scores were given in 5% steps.

The run-time per sample and per data set was significantly reduced approximately 10-fold by using the minRGB approach in comparison to the per-pixel classification methods. With the particularly used hardware configuration the image analysis time was up to 3-fold shorter than the time required for image acquisition, thus allowing implementation of image analysis in real-time.

## 4. Discussion

Powdery mildews possess a significant economic threat especially in the parts of the world with a warm and humid climate. The powdery mildews of wheat and barley PM are also an important model for studying plant-pathogen interactions. Therefore, this disease has long been in focus of the phenomics platform developers but the available tools are still relatively limited. Some methods are based on measuring the enzymatic activity of the infected tissue (Kuska et al. 2018), chlorophyll fluorescence (Brugger, Kuska, and Mahlein 2018) or on quantitative PCR of fungal genes (Wessling and Panstruga 2012) but more commonly optical sensors and computer vision approaches are used. Hyperspectral imaging is using the information about the reflectance of the tissues in a wide range of wavelengths and may visualize the disease symptoms in relatively early stages (Knauer et al. 2017, Thomas et al. 2018). The multispectral imaging is done with only a few but usually highly informative wavelengths thus greatly reducing the cost of equipment and the amount of raw data. However, the most common type of optical sensors is using the entire visible and a part of the near-visible spectrum. These sensors are either with integral wide band filter-matrices for limiting the sensitivity on pixel level to specific wavelengths (e.g. RGB cameras) or without wavelengths discrimination (grayscale cameras) but often with external filters and/or illumination sources with a discrete wavelength band. In our system, we have selected a CCD sensor of the second type (grayscale) to avoid some of the inbuilt problems of the RGB cameras (e.g. pixel values interpolation and lowered quantum efficiency). Instead of using filters for specific wavelengths we decided to use narrow bandwidth isotropic LED light sources, thus avoiding the use of motorized filter magazines and losing quantum efficiency. The nature of the samples (non-moving fixed objects) allows the acquiring of several images per object and combining the data without complicated merging methods. The leave samples were fixed in standardized containers (micro-titer plates), which greatly simplify the hardware design allowing the use of commercially available components such as the plate crane. The white plastic frames that keep the leaves fixed in the plates are at the same time used to define the area of interests, where the leaves are located.

Several machine learning and segmentation approaches were tested in order to find the most efficient algorithm for disease quantification. The most informative features for the machine learning approach were the H-, B- and a-channel of the resp. HSV, RGB and Lab color spaces. Among the tested texture features the Haralick was the by far most informative. A combined pixel classification based on color and texture features was tested as well but without significant improvement compared to the single features. Three different classifiers were tested and the Random forest (RF) performed slightly better than Support Vector Classifier (SCV) and much better than Linear SVC. Also, RF with a different number of trees was tested and the number of 50 trees was found to be optimal.

Astonishingly, among all tested segmentation approaches, the most accurate and efficient technique was the simple method of minimum RGB (minRGB). This filter was able to reliably detect the infected leaf area and to reduce the signal from disease-unrelated necrotic brown spots, which were of a particular problem in nearly all other approaches. Besides, the hardware workload and the calculation time for computing minRGB filter were significantly lower than of any other method. Finally, the minRGB was the segmentation method of choice, which was implemented into the image analysis pipeline.

The prediction results were validated by three other direct and indirect quantification methods – a manual scoring, as mean value of the scores of six different persons; quantitative PCR (qPCR); and inoculum density as number of spores per square millimetre of leaf surface. Although the genomic qPCR provides a nearly direct estimation of the total fungal biomass, it is a complex method, which is influenced by many factors, such as genomic DNA isolation and quality, primer design, PCR efficiency, detection sensitivity etc. Also, the measured quantity depends on both visible (on the leaf surface) and the invisible (too small or internal) fungal structures and is therefore not necessarily in perfect correlation to the visible disease symptoms. The inoculum density is rather an indirect parameter, which gives the infection pressure and the potential for the formation of fungal colonies, but the formation of the final fungal biomass depends on several other biotic and abiotic factors, as spore fitness and aggressivity, plant response and support of the fungal growth, temperature, humidity, etc. The mean scoring value of several persons provides a very robust parameter and therefore it was the method of choice for calibration of the automatic prediction.

### 4.1.1. Hardware and software assembly of the Macrophenomics platform

Based on the experimental results we have selected the best performing software protocols and combined them to a fully automated phenotyping pipeline. The software part runs both on front and back end in the Microsoft Windows 64-bit environment.

The Macrobot system itself is equipped with a custom imaging system software developed by Fraunhofer IFF. Several software modules control all actors and sensors in the system providing services to a service manager. The flow control for the imaging process is achieved by script programming, which enables a change in the imaging process without re-implementing the different software modules and makes extensions to the system easy and efficient. System modules providing a graphical user interface are organized in a reconfigurable user interface, which can be arranged to the needs of the system user without re-implementation. The imaging system generates a structured dataset for the subsequent image analysis.

The image analysis part is implemented on Windows 7 operating system and requires Python 2.7 or higher depending on the NumPy, opencv-python, scikit-learn and scikit-image open-source libraries.

The resulting pipeline provides precise phenomics data for the powdery mildew resistance trait in cereals. Exact, reproducible and non-biased phenotyping data are essential for discovering quantitative trait loci (QTL) with a minor but additive effect, which are contributing to a durable and broad-spectrum quantitative disease resistance. Although the manual evaluation of this phenotype is still the gold standard, the poor reproducibility frequently observed between the results of one assessing person to another and between the assessments is often insufficient to provide a solid statistical background for discovering minor resistance traits. In this work, we demonstrate that our Macrophenomics module can provide reliable and reproducible data in a very good correlation to the average score of multiple assessing persons and it can outperform single scoring persons by the accuracy of infection area estimation. The module is also fully open for adaptation to other than powdery mildew leaf diseases as such as different spot-, blight- and rust diseases caused by several fungal, viral and bacterial pathogens, such as yellow and brown rusts (*Puccinia sp.*), Septoria leaf blotch (*Zymoseptoria tritici*), spot blotch (*Bipolaris sorokiniana*), Bacterial leaf blight (*Pseudomonas syringae*), bacterial leaf streak and black chaff (*Xanthomonas translucens*), Barley yellow dwarf virus, etc. However, an important limitation is that the tested objects must fit into a standard MTP container (app. 12 x 8 x 1 cm), which includes samples like detached leaves, seeds, stem and root fragments, cereal spikes, and small whole plants.

## Acknowledgments

We would like to acknowledge the following members of the former Pathogen Stress Genomics group at IPK Gatersleben: G. Brantin, Dr W. Chen, Dr D. Nowara, and Dr J. Rajaraman for their contribution to the manual disease quantification data. Further thanks to Dr. D. Nowara for providing the primers and probes for the qPCR experiment.

## Author contributions

SL designed and programmed the image analysis software and performed the validation experiments; MS contributed in writing the paper and in developing the machine learning approach; ML performed the wheat infection experiment; FM, AB, DK and US designed and developed the Macrobot hardware and controlling software, PS developed the concept and acquired funding; AD contributed to writing the manuscript; DD contributed into designing of the image analysis software and hardware, performed the experiments and wrote the manuscript.

## Funding

This work was performed within the German Plant Phenotyping Network (DPPN) which is funded by the German Federal Ministry of Education and Research (BMBF) (project identification number: 031A053).

## Competing interests

The authors declare that there is no conflict of interest regarding the publication of this article.

## Data availability

Image data used for validation of the Macrobot algorithm is available at (Douchkov D 2019)

## 6. Supplemental figures and tables

**Supplemental figure S1.**
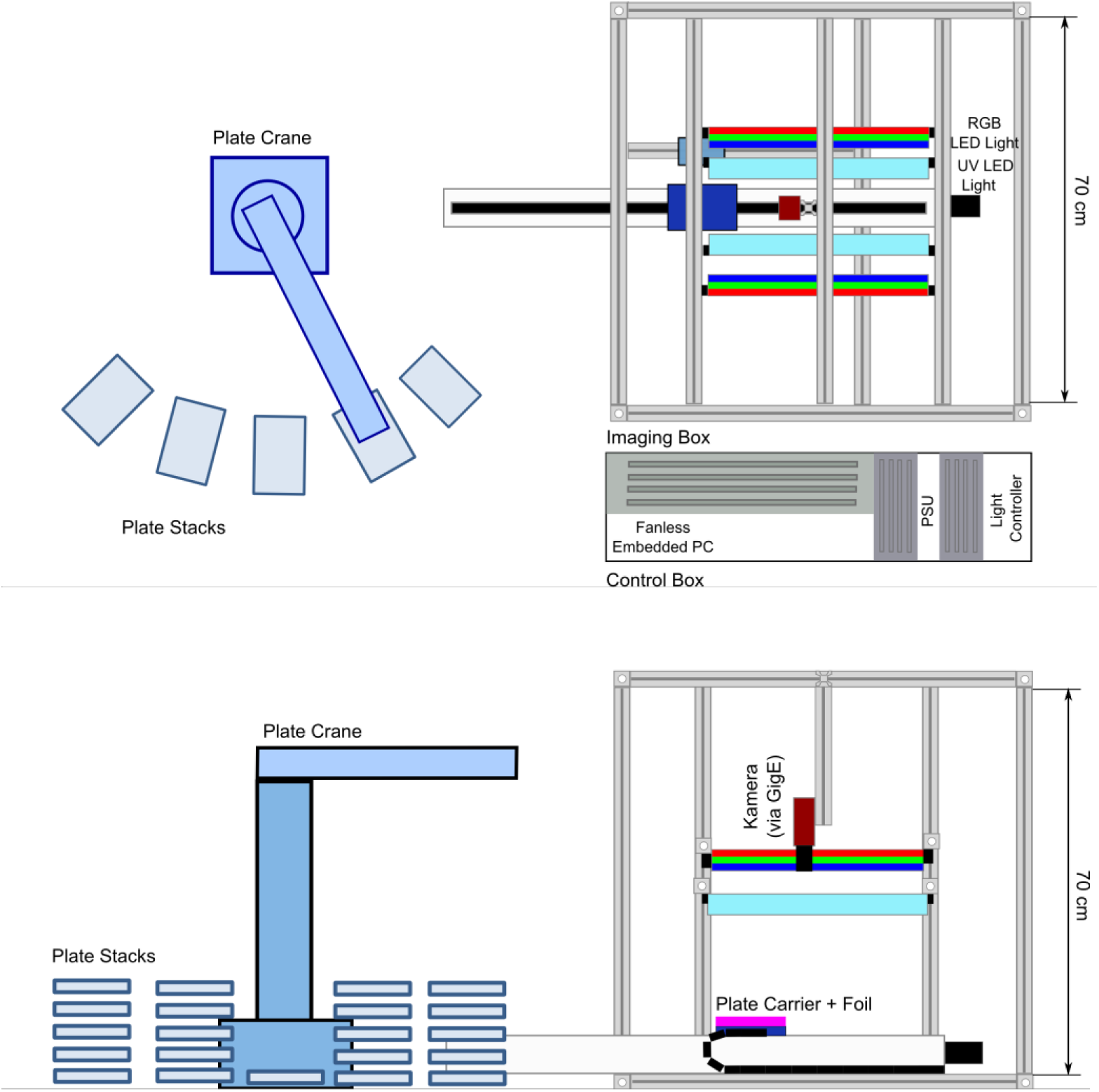
Schematic drawing of the image acquisition hardware of the Macrophenomics module Macrobot 2.0 (top and side views).

**Supplemental Table S1.** Pierson’s r^2^ coefficients of different manual and automated scorings (to **Figure 9**)

| Expert 1 | Expert 2 | Expert 3 | Expert 4 | Expert 5 | Expert 6 | Mean man. | qPCR | Inoc. dens. |  |
| --- | --- | --- | --- | --- | --- | --- | --- | --- | --- |
| 0.71 | 0.68 | 0.69 | 0.59 | 0.62 | 0.58 | 0.78 | 0.52 | 0.63 | Automated score |
|  | 0.71 | 0.79 | 0.60 | 0.60 | 0.54 | 0.88 | 0.66 | 0.60 | Expert 1 |
|  |  | 0.76 | 0.67 | 0.58 | 0.59 | 0.86 | 0.68 | 0.67 | Expert 2 |
|  |  |  | 0.64 | 0.63 | 0.65 | 0.91 | 0.74 | 0.62 | Expert 3 |
|  |  |  |  | 0.56 | 0.51 | 0.77 | 0.66 | 0.56 | Expert 4 |
|  |  |  |  |  | 0.57 | 0.78 | 0.57 | 0.60 | Expert 5 |
|  |  |  |  |  |  | 0.76 | 0.52 | 0.58 | Expert 6 |
|  |  |  |  |  |  |  | 0.77 | 0.74 | Mean man. |
|  |  |  |  |  |  |  |  | 0.56 | qPCR |

**Supplemental Table S2.**
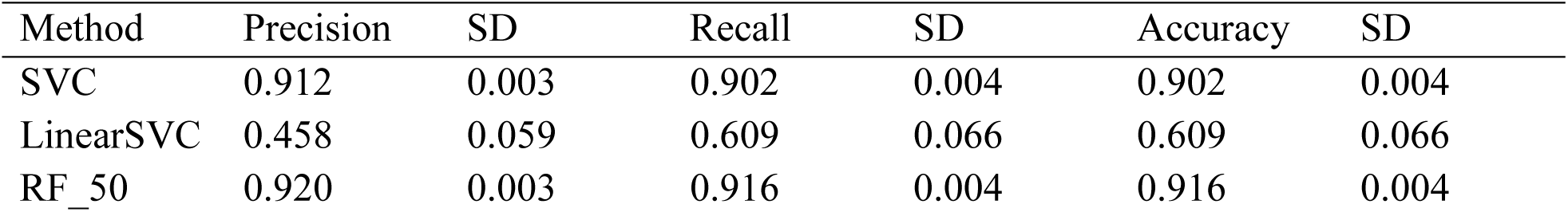
Values to **Figure 4A** - Evaluation of different classifiers on HSV_H_channel (5000 pixels per class, n=10).

**Supplemental Table S3.**
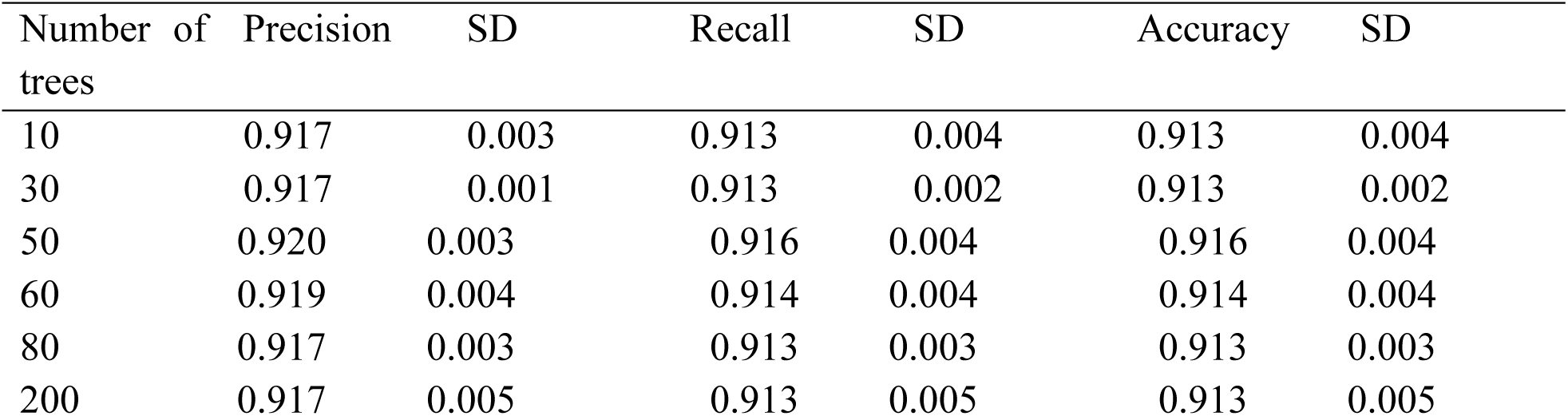
Values to **Figure 4B** - Evaluation of Random forest classifier with different number of trees on HSV_H_channel (5000 pixels per class, n=10).

**Supplemental Table S4.** Values to **Figure 5A** - Evaluation of different color pixel classification method (n=10)

| Method | Precision | SD | Recall | SD | Accuracy | SD |
| --- | --- | --- | --- | --- | --- | --- |
| RGB | 0.7632 | 0.0062 | 0.7577 | 0.0059 | 0.7577 | 0.0059 |
| RGB_B_channel | 0.8390 | 0.0041 | 0.8308 | 0.0046 | 0.8308 | 0.0046 |
| RGB_G_channel | 0.7355 | 0.0063 | 0.7374 | 0.0063 | 0.7374 | 0.0063 |
| RGB_R_channel | 0.6601 | 0.0058 | 0.6521 | 0.0065 | 0.6521 | 0.0065 |
| HSV_H_channel | 0.9189 | 0.0034 | 0.9140 | 0.0038 | 0.9140 | 0.0038 |
| HSV_S_channel | 0.6371 | 0.0068 | 0.6167 | 0.0056 | 0.6167 | 0.0056 |
| LAB_A_channel | 0.8750 | 0.0053 | 0.8605 | 0.0047 | 0.8605 | 0.0047 |
| LAB_B_channel | 0.7214 | 0.0050 | 0.7195 | 0.0051 | 0.7195 | 0.0051 |
| Grayscale | 0.7487 | 0.0071 | 0.7446 | 0.0055 | 0.7446 | 0.0055 |
| HSV | 0.6324 | 0.0044 | 0.6172 | 0.0039 | 0.6172 | 0.0039 |
| LAB | 0.7994 | 0.0020 | 0.7871 | 0.0031 | 0.7871 | 0.0031 |

**Supplemental Table S5.** Values to **Figure 5B** - Evaluation of texture features (n=10)

| Method | Precision | SD | Recall | SD | Accuracy | SD |
| --- | --- | --- | --- | --- | --- | --- |
| LBP | 0.1758 | 0.0159 | 0.4189 | 0.0190 | 0.4189 | 0.0190 |
| Haralick | 0.9836 | 0.0030 | 0.9522 | 0.0017 | 0.9746 | 0.0014 |
| PFTAS | 0.4950 | 0.0027 | 0.5202 | 0.0027 | 0.5202 | 0.0027 |

